# Dynamic changes of genomic and transcriptomic tumor diversity during melanoma progression

**DOI:** 10.1101/2021.03.18.435947

**Authors:** Ariane L. Moore, Christos Dimitrakopoulos, Stefanie Flückiger-Mangual, Lukas Frischknecht, Christian Beisel, Wilhelm Krek, Niko Beerenwinkel

**Affiliations:** Department of Biosystems Science and Engineering, ETH Zurich, Mattenstrasse 26, 4058 Basel, Switzerland; SIB Swiss Institute of Bioinformatics, Mattenstrasse 26, 4058 Basel, Switzerland; Institute for Molecular Health Sciences, ETH Zurich, Otto-Stern-Weg 7, 8093 Zurich, Switzerland

## Abstract

Metastatic melanoma is an aggressive disease that is notoriously difficult to treat. Here, we used a BRAF^V600E^-driven mouse model to study melanoma progression and metastasis in the absence of extrinsic mutagenic factors such as UV radiation. The tumors were grown for different time periods in seven different mice such that disease progression was tracked over time. We sequenced the whole exome and whole transcriptome of 16 melanoma samples from these mice. The timed samples revealed that many genes are increasingly upregulated during progression, such as genes involved in extracellular matrix organization, and *Vegfc*, a factor that induces lymphatic vessel growth and metastases to lymph nodes, which was identified as an important signaling hub in our analysis. The *Vegfc* gene interaction network highlighted multiple genes from the MAPK pathway that are increasingly deregulated during tumor evolution. The genomic diversity of primary and metastatic tumors showed an increase over time. Additionally, our integrative multi-omics analysis adds further evidence that *Rock1* is a crucial player driving metastasis formation, and may be upregulated due to mutations in the PI3K/AKT pathway. In summary, our study contributes towards an improved understanding of melanoma which may eventually lead to improved treatment strategies.

## Introduction

Metastatic melanoma is the most lethal type of skin cancer^1^. The main contributor to melanoma development is exposure of the skin to UV radiation in sunlight^1^. The incidence rate of melanoma in the United States and Europe has been rising over the past decades^2,3^. The estimated number of new melanoma cases and deaths in 2018 in the United States is 91,270 and 9,320, respectively^2^. The five-year survival rate of metastasized melanoma in the United States is estimated to be around 20%^2^. Progress in understanding the molecular underpinnings of this disease, as well as earlier detection has led to improved survival rates^1,3^. However, resistance emerges to both oncogene-targeting agents as well immunotherapies, which warrants the need for an improved understanding of tumor progression^1,4-6^.

Extensive research has led to the identification of several oncogenes, tumor suppressors and key pathways in melanoma that are frequently altered and present potential drug targets^7^. Most commonly, melanomas are driven by oncogenic MAPK signaling driven by activating mutations in *BRAF* or *NRAS* ^8-10^. Additionally, the inactivation of the tumor suppressor gene *PTEN* is a frequent event in melanomas^8,11^, which activates the PI3K/AKT pathway and contributes to tumor invasion^1^. Furthermore, Wnt signaling is often activated and contributes to cell proliferation^1^.

Dankort and colleagues introduced a mouse model of tamoxifen-inducible, *Braf*-driven and *Pten*-deficient melanoma that mimics the human disease^8^. It offers the possibility to analyze melanoma development and to characterize the genomic and transcriptomic alterations that accompany the mutations in *Braf* and *Pten*. For instance, it has been shown that *Ctnnb1*, which encodes for Catenin beta and belongs to the Wnt pathway, is another key player that is significantly mutated in melanoma^12^ and, when activated, drives tumor development together with *Braf* and *Pten* ^13^. This mouse model allows to study melanoma progression and metastasis without the large number of potential passenger mutations that may occur in UV-damaged skin and is usually observed in melanoma^14^.

Here, we apply the mouse model of *Pten*-deactivated and *Braf*-activated melanomas to study melanoma progression and metastasis in detail. In seven different mice, melanoma was induced on the mouse tail and the tumors were grown over different time periods in each mouse. As a result, the collected samples are from seven different time points of days after tumor initiation (Figure 1A, upper panel). From each mouse, a primary melanoma sample was collected. Additionally, from each but one mouse, one or two samples from metastases in the lymph nodes were extracted. One normal tail sample served as control. All samples were subjected to whole-exome sequencing (WES), and RNA sequencing (RNA-seq). The samples were collected based on time after tamoxifen-driven tumor induction and not based on tumor size. Hence, the experimental design that includes the timed samples allows the examination of the genomic and transcriptomic changes of melanoma progression over time (Figure 1a, lower panel). Additionally, the comparison of metastatic and primary tumor sites enables the investigation of tumor diversity over time. Moreover, we combined genomic and transcriptomic alterations through a network-based approach (Figure 1b) in order to interpret the set of molecular alterations from a systems perspective. Integrative network analysis can aid to understand the complex interactions between different molecules and to identify novel strategies to combat the disease^15,16^.

**Figure 1.**
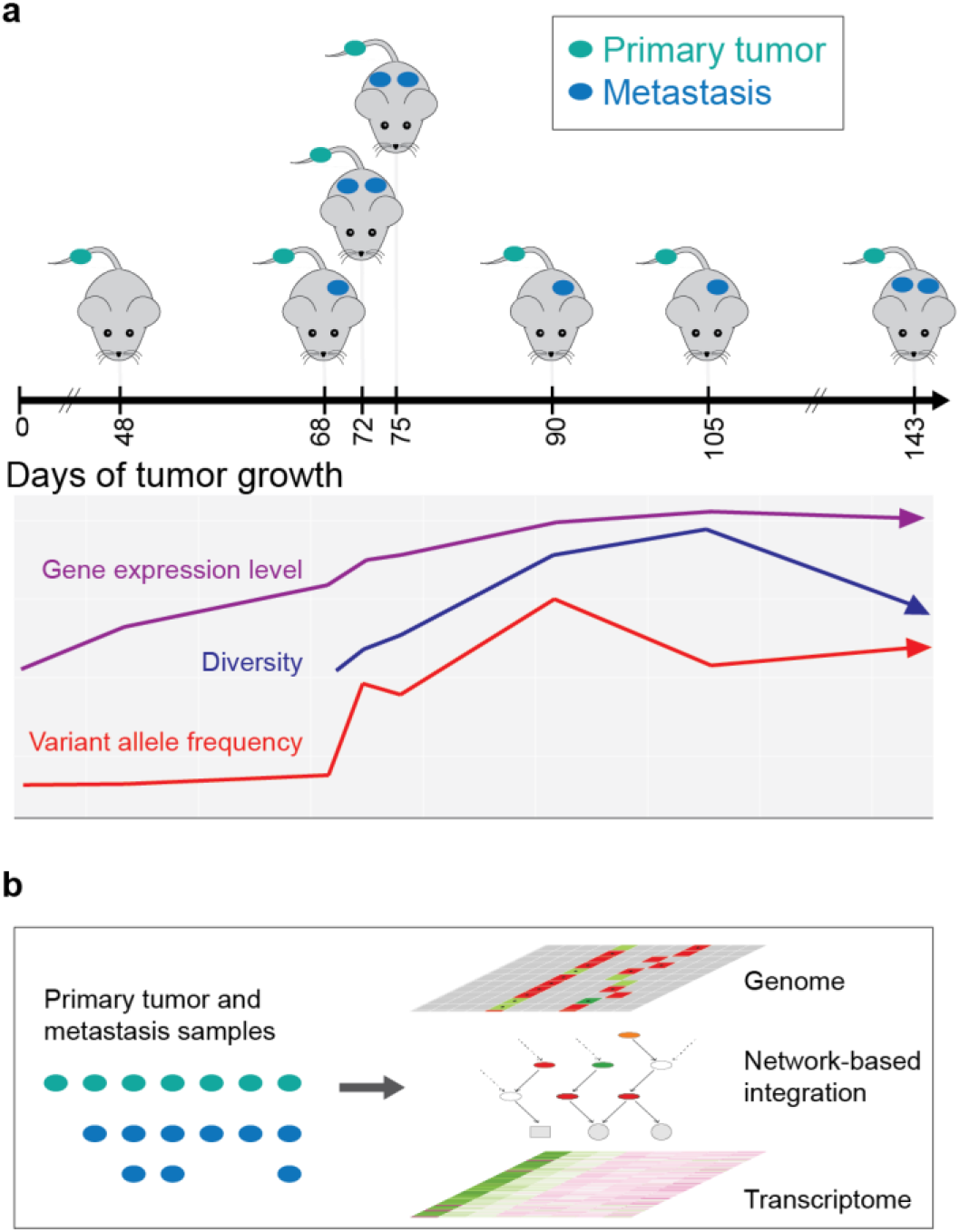
(a) Upper panel: Experimental design. The analysis includes a tamoxifen-inducible mouse model of *Braf*^V600E^-driven, *Pten*-deficient melanoma^8^ from seven time points of days of tumor growth. From each of the seven mice, one primary tumor sample was collected. For all but the first mouse, also one or two lymph node metastasis samples were extracted. As a control, a normal tail sample was collected. **Lower panel:** Schematic representation of the aspects under examination. The genomic and transcriptomic changes over time are investigated by tracking the variant allele frequency or gene expression level in the timed samples. Additionally, the comparison of primary and metastatic tumor sites enables an analysis of the diversity over time. **(b)** For each sample, whole-exome and RNA sequencing was performed, allowing network-based integration of genomic alterations and transcriptomic changes.

## Results

The collected samples are from seven different time points of tumor growth, namely 48, 68, 72, 75, 90, 105, and 143 days (Figure 1a, upper panel; Suppl. Table S1). The samples are referred to by the number of days of tumor growth, as well as the abbreviations “Tu” for primary tumor sample, and “Met” for metastasis samples (Suppl. Table S1). For instance, the primary tumor sample of the first mouse at 48 days of tumor growth is referred to by “48_Tu”. In the following, we will first provide an overview of the detected genomic changes. Next, the timed primary samples are in focus to highlight the transcriptomic changes over time, and network analysis is used to detect important signaling hubs that link genomic alterations to differentially expressed genes. Finally, the genomic and transcriptomic diversity is examined in detail to identify important aspects of metastasis formation.

### Overview of genomic changes

The average coverage in the WES samples was 65x (Suppl. Table S1). The median number of single-nucleotide variants (SNVs) and indels that were detected per sample was 25 (Materials and Methods). Overall, the number of genes affected by SNVs and indels was slightly lower for earlier samples, and higher for later samples (Figure 2a). However, this can be attributed mostly to two rather late metastasis samples that seem to be hypermutators (Figure 2a, upper panel; Suppl. Figure S1). These samples harbored many more mutations than all other samples, which affect known cancer genes (Figure 2a, lower panel). Among the mutated genes in the hypermutators, *Brca2, Pik3r1, Bclaf1, Chek2*, and *Rad51b* play a role in DNA repair and response to DNA damage^17-21^. This suggests that the DNA repair mechanism was defect leading to an accumulation of numerous SNVs and indels^22^. Moreover, the two hypermutators 75_Met2, and 105_Met have, with 45 and 13, the highest number of deleterious mutations compared to the other samples, which all have less than 10 variants classified as deleterious by SIFT^23^.

**Figure 2.**
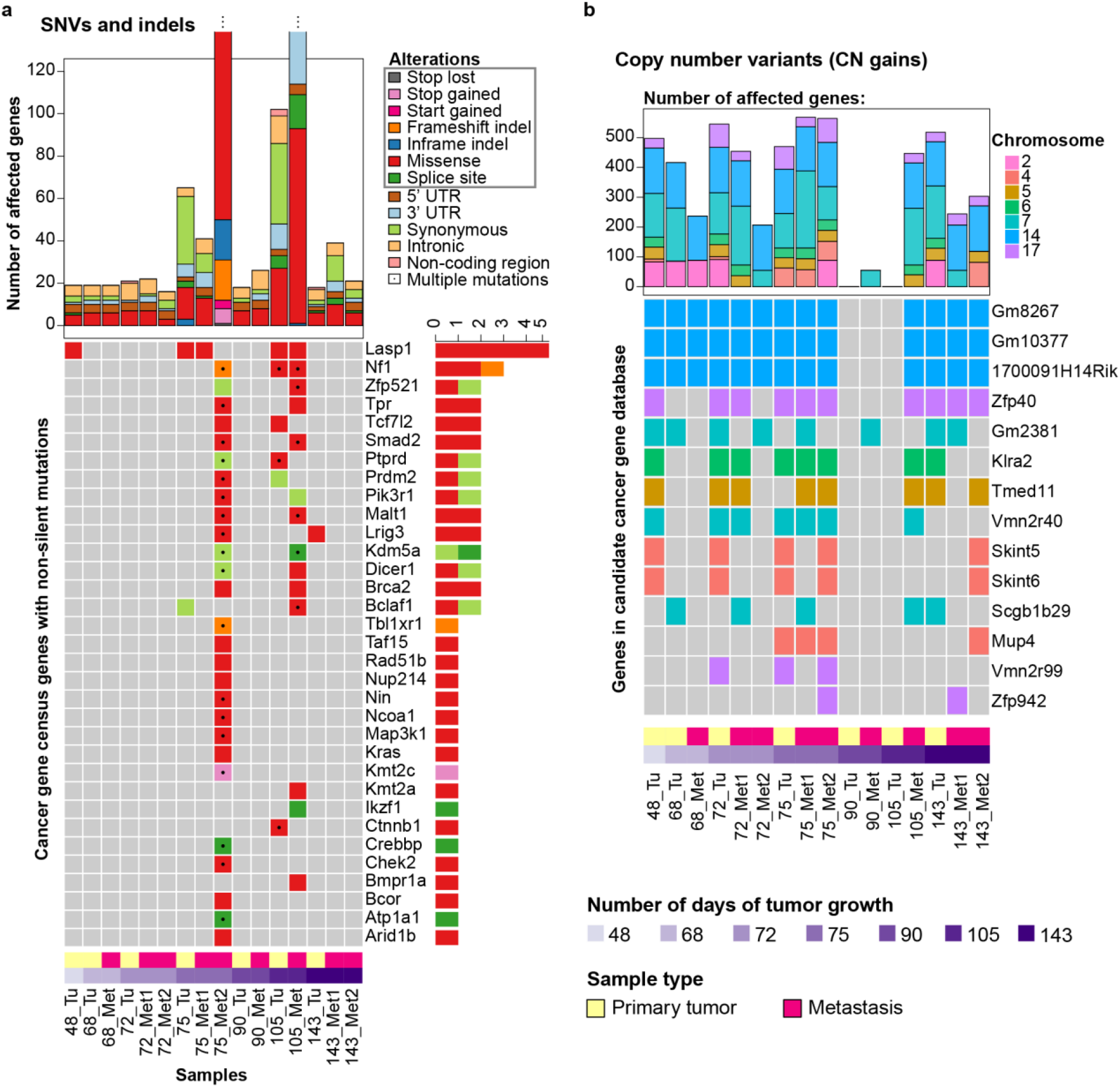
Overview of the genomic diversity. **(a)** Upper panel: The numbers of genes that are affected by a single-nucleotide variant (SNV) or indel per sample. The color indicates the annotation of the mutation (Full barplot in Suppl. Figure S1). Lower panel: The heatmap shows all cancer gene census genes that harbor a nonsynonymous mutation in a coding region or a splice site mutation (grey box in the legend ‘Alterations’). Multiple mutations in a gene are indicated with the black dot, and the color represents the annotation of highest priority, which is the same ordering as in the legend from top to bottom. For instance, if a gene had a missense and a synonymous mutation, the color will be red, and the black dot indicates that there were additional mutations. In order to clearly show which sample name is referring to which sample type and mouse, the numbers of days of tumor growth as well as the sample type is indicated. **(b)** Upper panel: The number of genes that are affected by a copy number change. The color indicates on which chromosome the copy number change occurred. The gonosomes are omitted from the plot and further analysis (Materials and Methods). All copy number events displayed were copy number gains. Lower panel: The heatmap depicting the genes affected by a copy number gain that are in the candidate cancer gene database^27^. As before, the numbers of days of tumor growth as well as the sample type is indicated for each sample.

In general, several of the mutated genes identified here have been associated with melanoma. *Nf1*, which was detected in three samples with a deleterious mutation^23^, is a known tumor suppressor gene and among the most commonly mutated genes in melanomas^24^. It was reported that melanomas with *NF1* mutations are linked to a higher mutational burden and shorter survival^12^. Furthermore, in combination with *BRAF* mutations, the inactivation of *NF1* has been shown to be a resistance mechanism against *BRAF* inhibitors^25^. Additionally, the identified genes *Crebbp, Ptprd, Map3k1* and *Bclaf1* are recurrently mutated in melanoma, and *Crebbp* was suggested to be a potential driver gene^14^. Also, mutations in *RAS* genes, such as *KRAS*, as well as mutations in the PI3K pathway, for instance, in *PIK3R1*, are frequently observed in melanoma^12^. By contrast, *LASP1* is mutated in less than 1% of TCGA melanoma samples^26^, but in our data, *Lasp1* is affected by a deleterious missense mutation in five of 16 samples.

The analysis of copy number variants (CNVs) (Materials and Methods) identified on average five copy number events per sample. These affected on average 345 genes per sample (Figure 2b, upper panel), and all copy number events were copy number gains. Among the affected genes, 14 are in the candidate cancer gene database (Figure 2b, lower panel), which comprises genes that are potential cancer driver genes in mice^27^. These include two genes encoding for Zinc finger proteins, *Zfp942* and *Zfp40*, which play a role in the regulation of transcription, as well as two genes involved in G-protein coupled receptor activity, namely *Vmn2r40* and *Vmn2r99* ^28^.

### Analysis of transcriptomic changes in the primary tumor samples over time

The experimental design allows analyzing the transcriptomic changes over time. First, the timed primary tumor samples were examined in detail. The principal component analysis (PCA) of the primary tumor samples and the normal sample revealed that the tumor samples become more different from the normal with increasing numbers of days of tumor growth (Figure 3a). In particular, the first two tumor samples, 48_Tu, and 68_Tu, are much closer to the normal than the later samples. The PCA of the primary tumor samples without the normal sample is similar to Figure 3a (Suppl. Figure S2), also supporting the clear indication that primary tumors over time become more different from the first time point. In order to detect differentially expressed genes, two comparisons were made with the primary samples and the normal sample. That is, in one comparison, all primary tumor samples were compared against the normal sample (“primary versus normal”), and in the other comparison, the timed primary tumor samples were divided into early and late to identify genes that become up- or downregulated during melanoma development (“primary late versus early”). The cutoff to divide the primary tumor samples into early versus late was 70 days of tumor growth, because this represented the major divide in the PCA (Figure 3a). Hence the early primary tumor samples are 48_Tu and 68_Tu, and the late primary tumor samples are 72_Tu, 75_Tu, 90_Tu, 105_Tu and 143_Tu.

**Figure 3.**
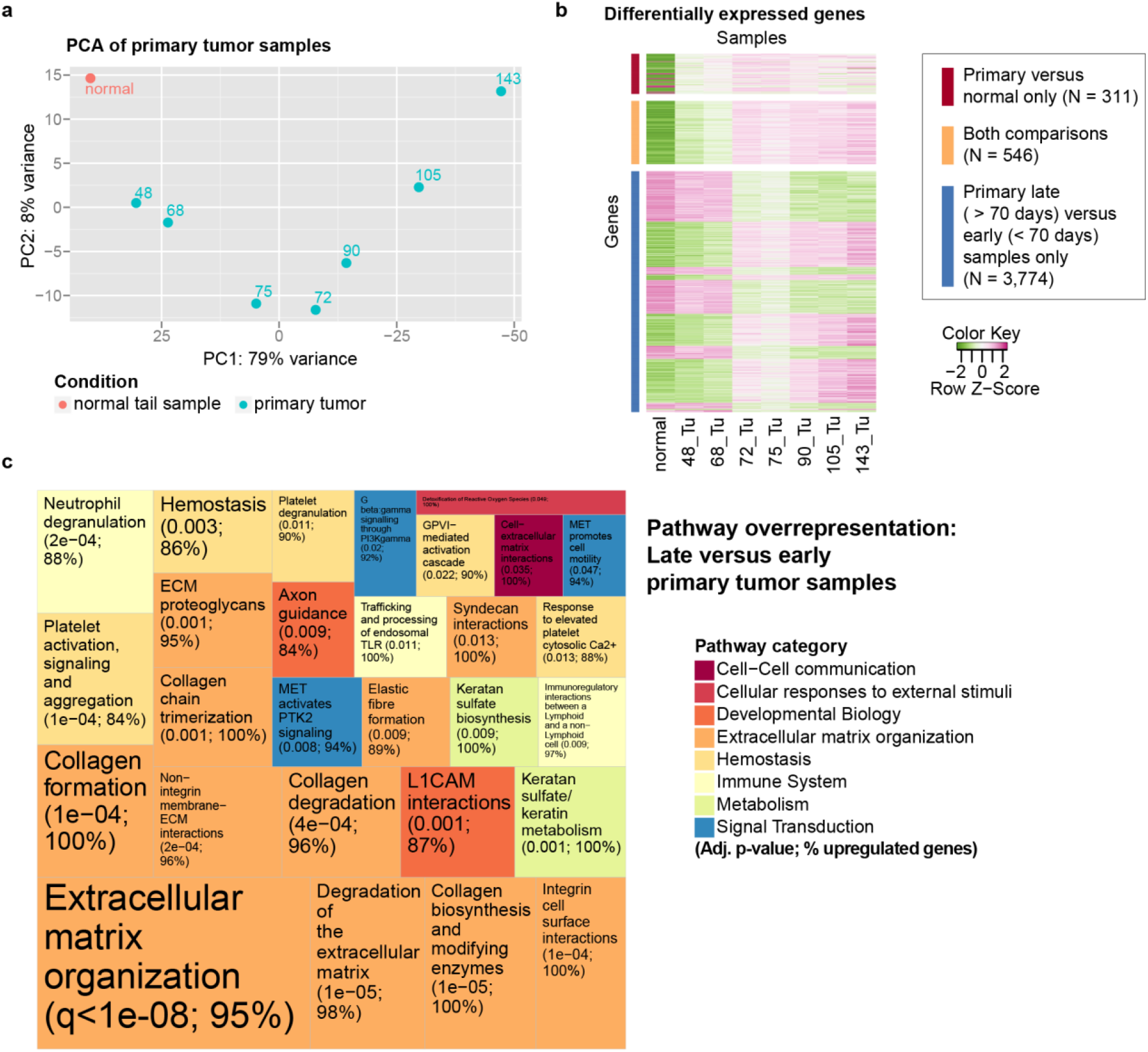
Transcriptome analysis of timed primary tumor samples. **(a)** Principal component analysis (PCA) of the expression levels of the 500 most variable genes when including all primary tumor samples and the normal sample. The number of days of tumor growth is indicated in the plot. **(b)** Differential gene expression analysis revealed 857 deregulated genes when comparing the primary samples to the normal. Most of them are upregulated as shown by the heatmap, where pink indicates higher expression levels, and green lower expression levels. The expression levels visualized here are normalized, and additionally centered and scaled such that each row has mean zero and standard deviation one. Moreover, differential gene expression analysis was performed to compare the five late primary tumor samples (> 70 days) to the two early primary tumor samples (48_Tu, 68_Tu). A total of 4,320 genes was differentially expressed in this comparison, where 546 genes were significantly upregulated in both comparisons. **(c)** Pathway overrepresentation analysis was performed with all 4,320 differentially expressed genes of the late versus early primary tumor samples comparison. The colors represent the top-level pathways of the Reactome hierarchy, and each individual pathway which was significantly overrepresented, is indicated in the treemap. The area of the tiles mirrors the significance level. The rounded adjusted p-value as well as the percentage of upregulated genes among the differentially expressed genes is indicated.

Differential gene expression analysis identified 857 deregulated genes when comparing all seven primary tumor samples to the normal sample (Figure 3b). The comparison of the five late primary samples versus the two early primary tumor samples resulted in 4,320 differentially expressed genes. A total of 546 genes are shared between these two comparisons (Figure 3b), and they are all upregulated genes that generally increase in expression level with increasing numbers of days of tumor growth. In order to identify the functional roles of the deregulated genes, overrepresentation analysis was performed. Among the differentially expressed genes of the comparison “primary late versus early”, those in pathways related to extracellular matrix organization were overrepresented and these genes were mostly upregulated (Figure 3c; Suppl. Figure S3). Degrading and remodeling of the extracellular matrix is important to create a tumor microenvironment that is permissive for invasion and metastasis^29,30^. The upregulation of genes involved in the extracellular matrix organization has been reported previously in melanoma and was linked to resistance to anti-PD-1 immunotherapy^31^. The overrepresented pathways among the deregulated genes of the “primary versus normal” comparison are in general similar to the ones from the “primary late versus early” comparison (Suppl. Figure S4), but the cell-extracellular matrix interactions seem to be a phenomenon that emerges rather late during tumor development (Figure 3c; Suppl. Figure S3).

### Network-based integration of genomic and transcriptomic changes over time

In order to jointly investigate the genomic and transcriptomic changes over time, the tool NetICS (Network-based Integration of Multi-omics Data)^15^ was applied. It integrates the genomic and transcriptomic changes through diffusion in a directed functional protein interaction network^15^. The underlying rationale is that differential gene expression may be explained by upstream mutated genes. NetICS diffuses the information from genomic alterations forward in the network, and transcriptomic changes are diffused backwards^15^. In this way, NetICS identifies major signaling hubs, so-called mediator genes that are relatively close in the network to upstream genomic alterations and downstream differentially expressed genes, and can be regarded as signal transducers that may propagate perturbations from mutations to differentially expressed genes^15^. Thereby, NetICS also tries to explain different genomic changes in the different samples that converge through a mediator to the same expression change. Here, the differentially expressed genes from the “primary versus normal” comparison were used, as well as genomic aberrations from all primary tumor samples. As a result, 42 mediator genes were identified (Suppl. Table S2) and ranked across all samples^15^. Among these, 27 genes were found in the candidate cancer gene database^27^, including the gene *Vegfc*. This gene was among the most highly ranked mediator genes, while additionally being highly significant (adjusted p-value < 10^−3^, permutation test). Therefore, *Vegfc* appears to be an important signaling hub between the mutated genes and the differentially expressed genes. The protein interaction network of *Vegfc* was analyzed including the direct downstream genes of *Vegfc* that are differentially expressed, as well as upstream genes with genomic alterations that are up to four levels upwards of *Vegfc* (Figure 4).

**Figure 4.**
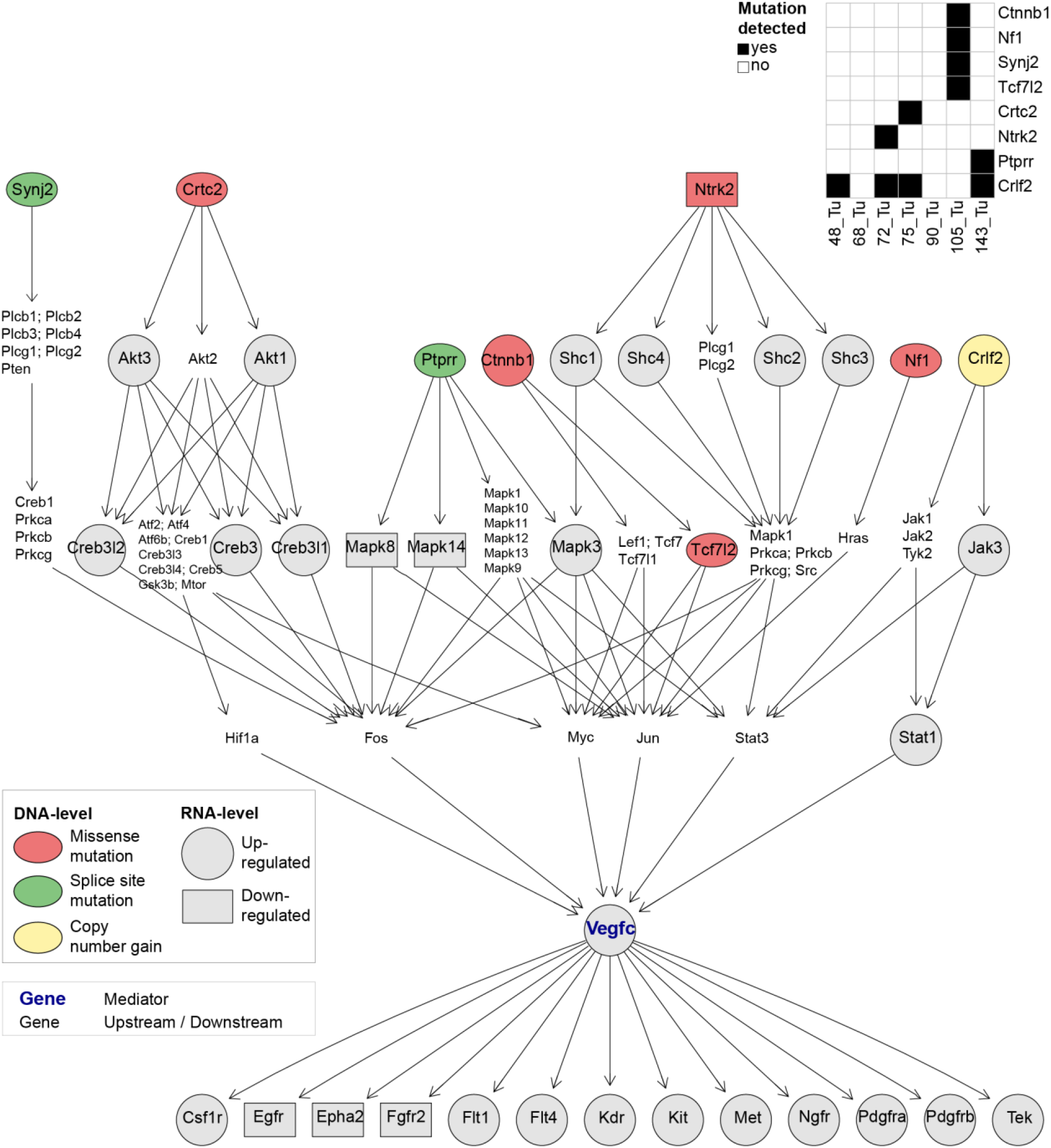
Network of mediator gene *Vegfc*. NetICS was performed to integrate RNA-seq and WES data. The displayed *Vegfc* network includes the upstream genes with non-silent genomic variation in at least one of the samples up to four levels upstream, as well as the direct downstream genes that are differentially expressed. Also, the genes connecting the upstream mutated genes with the mediator are indicated. Here, for reasons of biological interpretation, the deregulation of a gene is displayed if it is differentially expressed in either the “primary versus normal”, or the “primary late versus early” comparison. Additionally, for biological interpretation, also differential expression upstream of *Vegfc* is indicated, even if the upstream deregulated genes did not contribute to the mediator *Vegfc* in NetICS, since deregulation is diffused backwards. Some of the nodes summarize several genes in order to preserve space in the display. In such cases, the edges between nodes indicate that the edge is true for at least one of the genes in the summarized set. The table on the upper right indicates in which samples the mutations were detected.

In order to gain insight into the timing of these alterations and to determine which parts of the network are perturbed from the early beginning of neoplastic growth, and which changes occur later during the course of melanoma development, the genomic and transcriptomic alterations in this signaling network were examined in more detail. Pertaining to the genomic alterations, with the exception of the copy number gain in *Crlf2*, all mutations in the upstream genes of this *Vegfc* network occurred only in the late samples (Figure 5a, 5b). On the transcriptomic level, most changes occur when comparing the late to the early samples, too. More precisely, all but two of the significantly differentially expressed genes in this network occurred additionally, or exclusively, when comparing expression levels of late versus early primary tumor samples (Figure 5c). In particular, the expression level of *Vegfc* was significantly upregulated in both comparisons “primary versus normal” and “primary late versus early” (Figure 5c) indicating that the expression level of *Vegfc* is upregulated in all primary tumor samples, and that this trend continues during tumor evolution. This is in line with previous studies demonstrating that the expression level of *VEGFC* increases with melanoma progression^32^. More precisely, Goydos and Gorski examined VEGFC expression levels in 54 human melanoma samples from different phases of progression. They found that VEGFC expression was significantly higher in samples from later stages of progression^32^. Melanoma metastasizes to the lymph nodes through the lymphatic vessels^32^. *VEGFC* is a growth factor that stimulates lymphangiogenesis and hence is an important factor that enables metastasis through the lymphatic system^33^. It has been shown that overexpression of *VEGFC* is a predictor of the occurrence of lymph node metastases in melanoma^34,35^ and that it contributes to immune evasion^36^.

**Figure 5.**
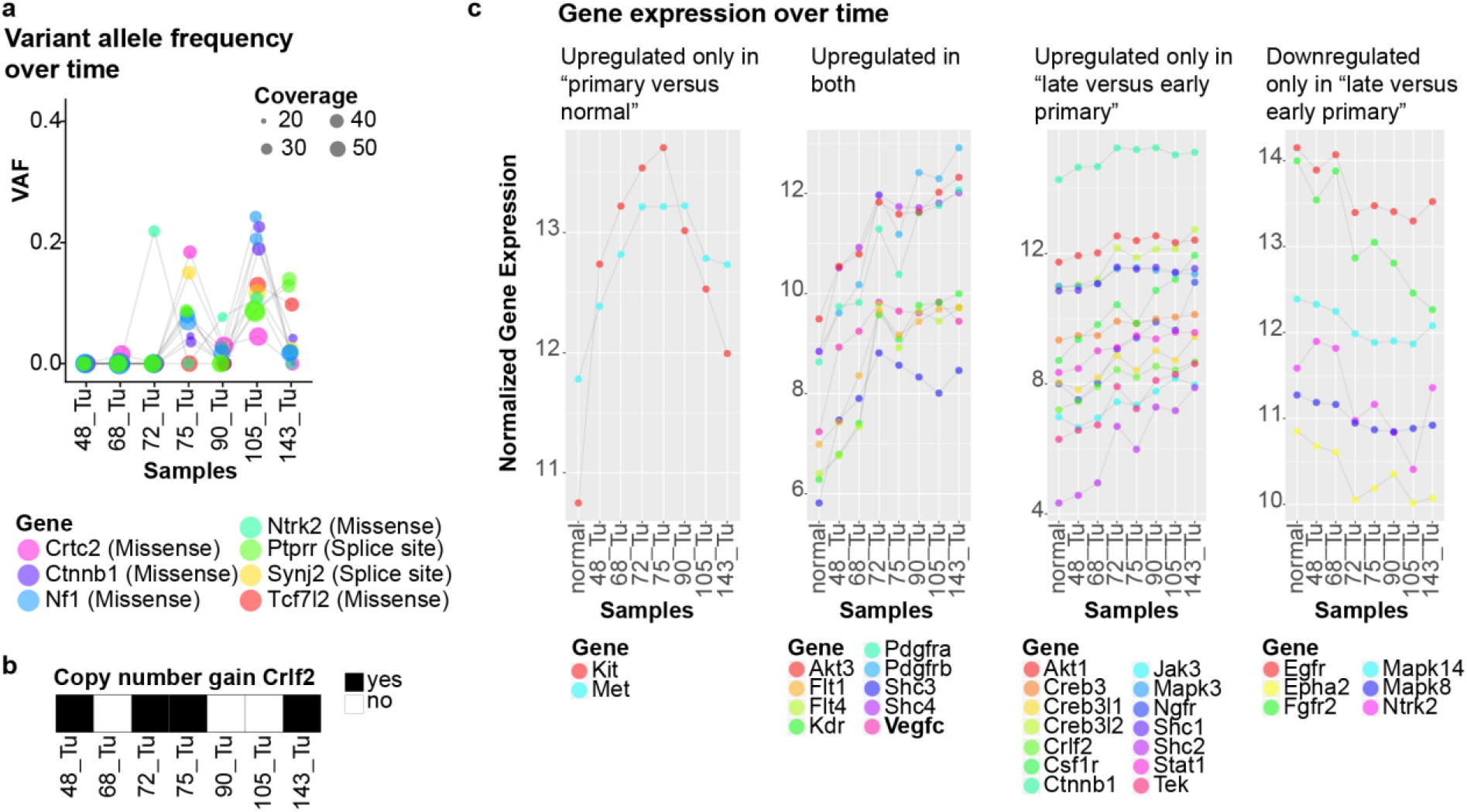
Genomic and transcriptomic alterations in the *Vegfc* mediator network. **(a)** Variant allele frequency of the upstream mutations in the primary tumor samples that are highlighted in the mediator network of Figure 4. **(b)** Copy number status of *Crlf2*. **(c)** Normalized gene expression levels of the differentially expressed genes in the network of Figure 4. In particular, the genes are separated into different categories depending on whether their deregulation occurred in all primary tumor samples, or only in the late samples. Genes with adjusted p-value < 0.01 were declared as being differentially expressed.

The upstream mutations of the *Vegfc* network include well known melanoma genes such as *Ctnnb1* and *Nf1*, as well as other, less common genes. *TCF7L2* encodes for TCF4, is part of the Wnt signaling pathway and is known to form a complex with *CTNNB1* ^37,38^. Mutations in this gene have been frequently observed in colorectal cancer^39^, whereas in melanoma, only 3% of TCGA samples show a *TCF7L2* mutation^26^. *NTRK2* is mutated in 5% of TCGA melanoma samples^26^. The missense mutation in *Ntrk2* detected here was classified as deleterious according to the SIFT prediction^23^. It was detected in sample 72_Tu, but can also be observed in 90_Tu, and 105_Tu with a variant allele frequency (VAF) of 8% and 11%, respectively, when considering all sequence reads at that position (Figure 5a). In the transcriptome, the expression level of *Ntrk2* is most downregulated in the three samples that show this mutation (Figure 5c), suggesting that the mutation is the cause of the downregulation. The known melanoma gene *Nf1* was affected by two deleterious missense mutations according to the SIFT prediction^23^ that were detected in the sample 105_Tu, but the samples 75_Tu, 90_Tu, and 143_Tu also show evidence in the sequence reads for these mutations with VAFs between 2-8% (Figure 5a).

Downstream of Vegfc, the differential gene expression analysis identified two growth factor receptors, *Egfr* and *Fgfr2* that are being downregulated in the course of tumor progression (Figure 5c). Two genes, *Kit* and *Met*, are upregulated in all primary tumor samples as compared to the normal, but their gene expression levels show a reversing trend in the latest tumor samples, which show decreased expression after 90 and 100 days of tumor growth (Figure 5c). *Kit* and *MET* are both known oncogenes that are overexpressed in some tumors^40,41^. *Pdgfra* and *Pdgfrb* are also upregulated in all primary tumor samples and their expression level increases in the late samples (Figure 5c). Overexpression of these two growth factor receptors has been documented for melanomas and other cancer types and was associated with cell proliferation and angiogenesis^42,43^. We also found that the receptor of Vegfc, Flt4, which is directly downstream of Vegfc in the network (Figure 5c), is increasingly upregulated during melanoma progression. Su and colleagues reviewed how VEGFC and its receptor Flt4, contribute to cancer progression, by, for instance, regulating invasion and cell proliferation^44^. Also, the expression level of VEGFC and Flt4 was linked to cancer progression in cervical cancer ^45^ and prostate cancer ^46^.

Overall, we observe that many of the genes in this *Vegfc* network are members of the MAPK signaling pathway, including *Vegfc* itself^47^. Precisely, besides *Vegfc*, eight of the upstream mutated or differentially expressed genes (*Ptprr, Ntrk2, Nf1, Akt1, Akt3, Mapk8, Mapk14, Mapk3*), and all of the 13 downstream deregulated genes are in the MAPK signaling pathway^47^. Our analysis demonstrated that most of these genomic and transcriptomic changes occur later during tumor development, or become stronger in the course of tumor evolution, as, for instance, the upregulation of *Vegfc*. This is in line with a recent melanoma study that also reported accumulating mutations and increased activity of the MAPK signaling pathway during tumor evolution^48-50^.

### Genomic diversity

For all but the first mouse, one or two metastasis samples from the lymph nodes have been extracted in addition to the primary tumor, allowing for the analysis of diversity within each mouse. Here, we examine the genomic diversity with respect to the time measured in days of tumor growth. In order to quantify the diversity and to compare it between the mice, we consider the measure of diversity introduced by Maley and colleagues^51^ and inspired by the taxonomic distinctness index in ecology^52^. It was originally defined as the fraction of loci that differ by LOH between two samples^51^. In order to make it independent of the number of samples from a neoplasm, Maley et al. used the mean pairwise genetic divergence in case there were more than two samples of a neoplasm^51^. They also showed that this measure is positively correlated with probability of progression from neoplasm to adenocarcinoma^51^. Here, we adopt the measure suggested by Maley and colleagues and consider instead of numbers of loci with LOH the numbers of genes that are affected by genomic alteration. Hence, we define the mean pairwise diversity score as

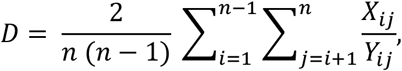

where *n* is the number of samples per individual mouse, *X*_*ij*_ is the number of genes that are affected by SNVs, indels, or copy number change in either sample *i* or *j* (but not both), and *Y*_*ij*_ is the total number of genes affected in both sample *i* and *j*. In other words, *D* is the mean fraction of privately mutated genes between all pairs of samples per mouse. The number of samples *n* per mouse can be different, but the mean pairwise diversity score is comparable between the mice, since the score is normalized by the number of sample pairs per mouse.

This measure was computed for all mice for which two or three samples were available. The analysis revealed that the diversity tends to increase with time of tumor evolution (Figure 6). The regression showed a significant short term increase in diversity between 68 and 105 days of tumor growth (p-value = 0.013; Suppl. Figure S5). The diversity score for the mouse at 75 days of tumor growth was reduced from 0.685 to 0.507 when excluding the hypermutator 75_Met2 (Suppl. Figure S5). However, the regression when excluding both hypermutators (75_Met2, 105_Met) still showed a significant short term increase in diversity between 68 and 90 days of tumor growth (p-value = 0.074; Suppl. Figure S5).

**Figure 6.**
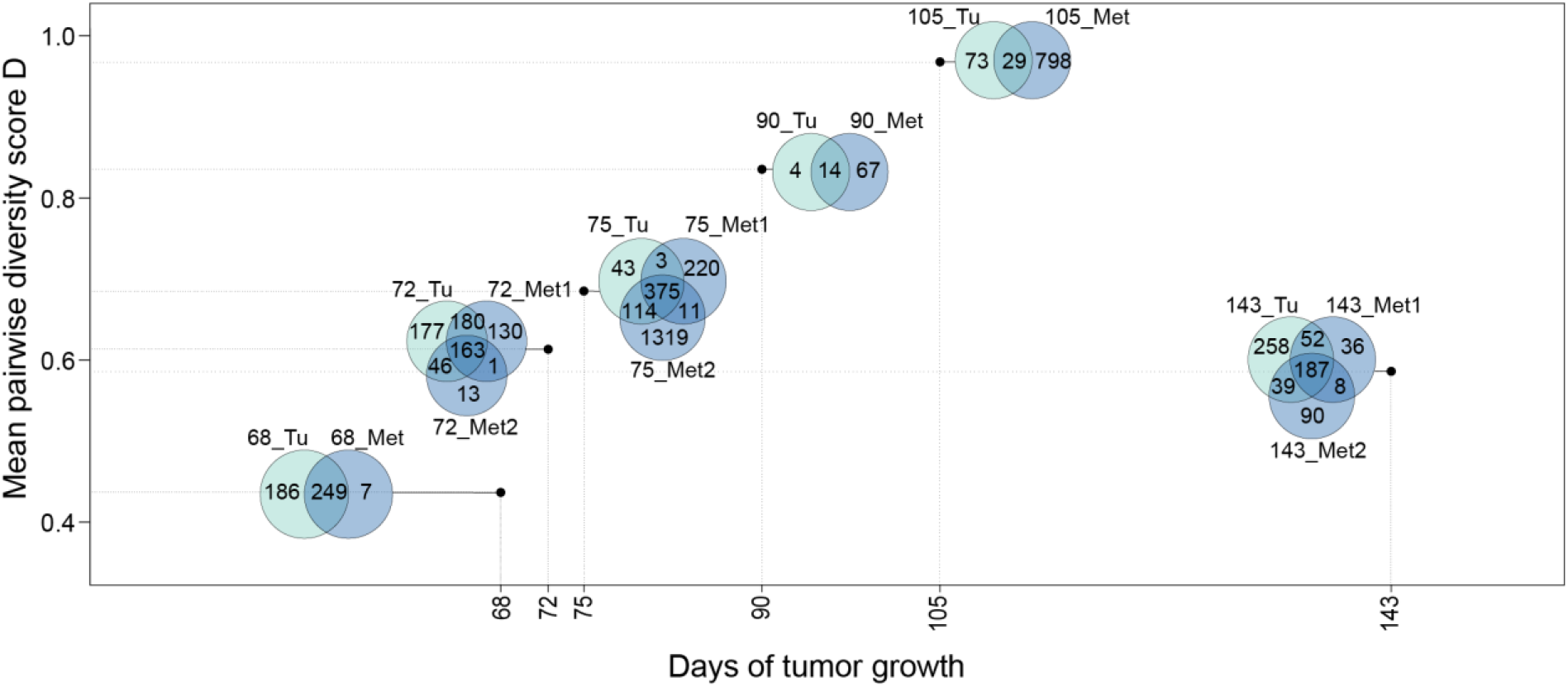
Genomic diversity over time. Venn diagrams displaying the number of genes affected by SNVs, indels or copy number change that are shared or private between the samples from each mice. The diversity score (y axis) is the mean fraction of privately mutated genes between all pairs of samples per mouse. In general, the diversity score tends to increase with the days of tumor growth. This trend is true for all but the last mouse, in which the tumor was grown much longer. The overall trend of increasing genomic diversity over time is still true when excluding the two hypermutators 75_Met2 and 105_Met (Suppl. Figure S5). Moreover, when considering the mice between 68 and 105 days of tumor growth, a significant short term increase in diversity can be observed (Suppl. Figure S5).

Additionally, phylogenetic trees can be used to interpret genomic diversity since they reveal the different subclones in tumors, including their genotypes and their evolutionary relationship^53^. This can be of crucial clinical importance, since, for instance, it has been shown that the order of mutations that occurred in tumors can influence factors such as treatment response^54^. Here, the phylogenetic trees were inferred using Cloe^55^ and most trees were linear (Suppl. Figure S6). This finding is in line with those of McGranahan and Swanton, who summarized the phylogenetic trees of different cancer types, and showed that evolutionary trees from melanoma are mostly linear with the bulk of the mutations being clonal^56^.

### Analysis of transcriptomic changes between the metastasis and the primary tumor samples

In order to identify which genes or pathways are deregulated in the metastasis samples as compared to the primary samples or the normal, the RNA-seq data from all samples were examined. PCA revealed that the primary tumor samples and metastasis samples differ quite substantially on the transcriptomic level (Figure 7a), albeit part of these differences may be attributable to the fact that the metastasis samples were extracted from the lymph nodes, whereas the primary tumor samples were from the tail. The comparison of all nine metastatic samples versus the normal sample (“metastasis versus normal”) resulted in 11,001 deregulated genes (Figure 7b). Differential gene expression analysis between all nine metastases and the 7 primary tumor samples (“metastasis versus primary tumor samples”) revealed 14,999 differentially expressed genes. A total of 8,900 genes are shared between both comparisons. Overall, half of the genes are upregulated, and the other half downregulated. Pathway overrepresentation analysis was performed to map the functions of the deregulated genes. Compared to the primary tumor samples, the genes involved in extracellular matrix organization are not upregulated any more in the metastases (Figure 7c; Suppl. Figure S3; Suppl. Figure S7). Instead, the deregulation of genes in nonsense-mediated decay is significant, and 90% of genes in this process are upregulated (Figure 7c). This can also be observed when comparing the metastasis samples to the normal sample (Suppl. Figure S7). The overexpression of genes involved in nonsense-mediated decay has been reported in cancer^57,58^. It is a process by which mutated, and also intact, transcripts are being degraded, and hence fulfils a crucial function in regulating the expression levels of other genes^57,58^. Furthermore, genes involved in keratinization are significantly downregulated compared to the primary tumor samples (Figure 7c). Keratins have been associated with cancer cell invasion and metastasis^59^. While metastasis samples do resemble their tissue of origin^60^, these aforementioned differences between the metastasis and primary tumor samples should be interpreted with caution, as they could also stem from tissue-related differences.

**Figure 7.**
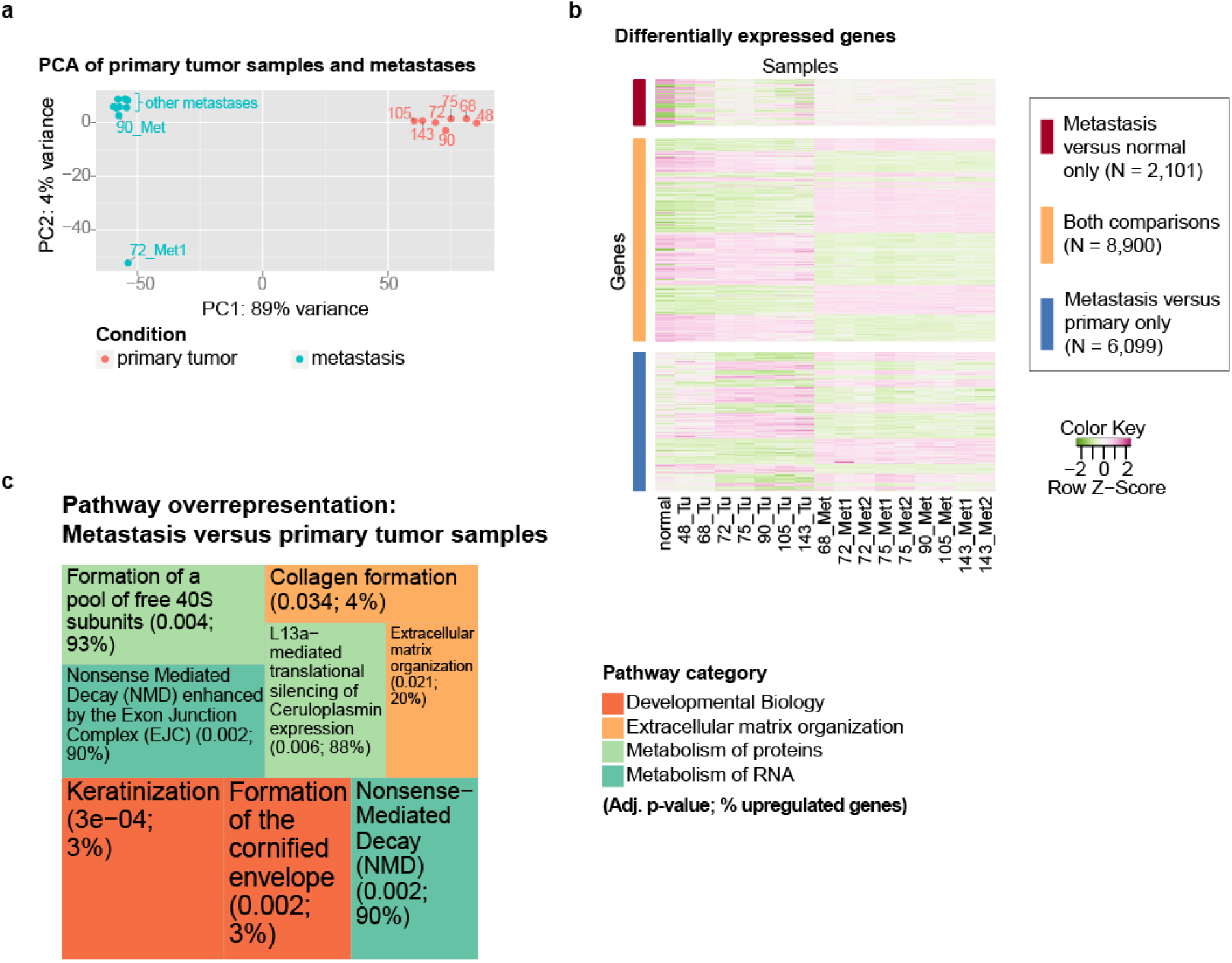
Transcriptomes of metastasis samples in comparison to primary tumor samples.**(a)** Principal component analysis (PCA) of expression levels of the 500 most variable genes when including all metastasis and primary tumor samples. The number of days of tumor growth is indicated in the plot. **(b)** Differential gene expression analysis revealed 11,001 deregulated genes when comparing the metastasis samples to the normal. Additionally, differential gene expression analysis was performed to compare the metastasis samples to the primary tumor samples. A total of 14,999 genes was differentially expressed in this comparison, where the 8,900 genes were significantly upregulated in both comparisons. **(c)** Pathway overrepresentation analysis was performed with all differentially expressed genes of the metastasis versus primary comparison. The colors represent the top-level pathways of the Reactome hierarchy, and each individual pathway which was significantly overrepresented, is indicated in the treemap. The area of the tiles mirrors the significance level.

### Network-based integration of genomic and transcriptomic changes between the metastasis and primary tumor samples

As before, NetICS was used to combine the mutations and differential gene expression patterns of primary and metastatic samples through diffusion in a directed functional protein interaction network^15^. Here, all differentially expressed genes from the “metastasis versus primary tumor samples” were considered, as well as mutated genes that have been detected in either the primary, or the metastatic samples. NetICS identified four significant mediator genes (Suppl. Table S3), of which three (*Tsc1, Rock1, Spred1*) are also candidate cancer genes^27^. *Rock1* (Rho-associated protein kinase 1) was selected for a more detailed analysis of its direct network, because it was highly significant (adjusted p-value < 10^−3^, permutation test). This network analysis included the direct downstream genes of *Rock1* that are differentially expressed, as well as upstream genes with genomic alterations that are up to four levels upwards from *Rock1* (Figure 8). Many of the upstream mutations were detected in the two metastasis samples with very high mutational burden, but also four other samples contribute to the set of upstream mutated genes (Figure 8).

**Figure 8.**
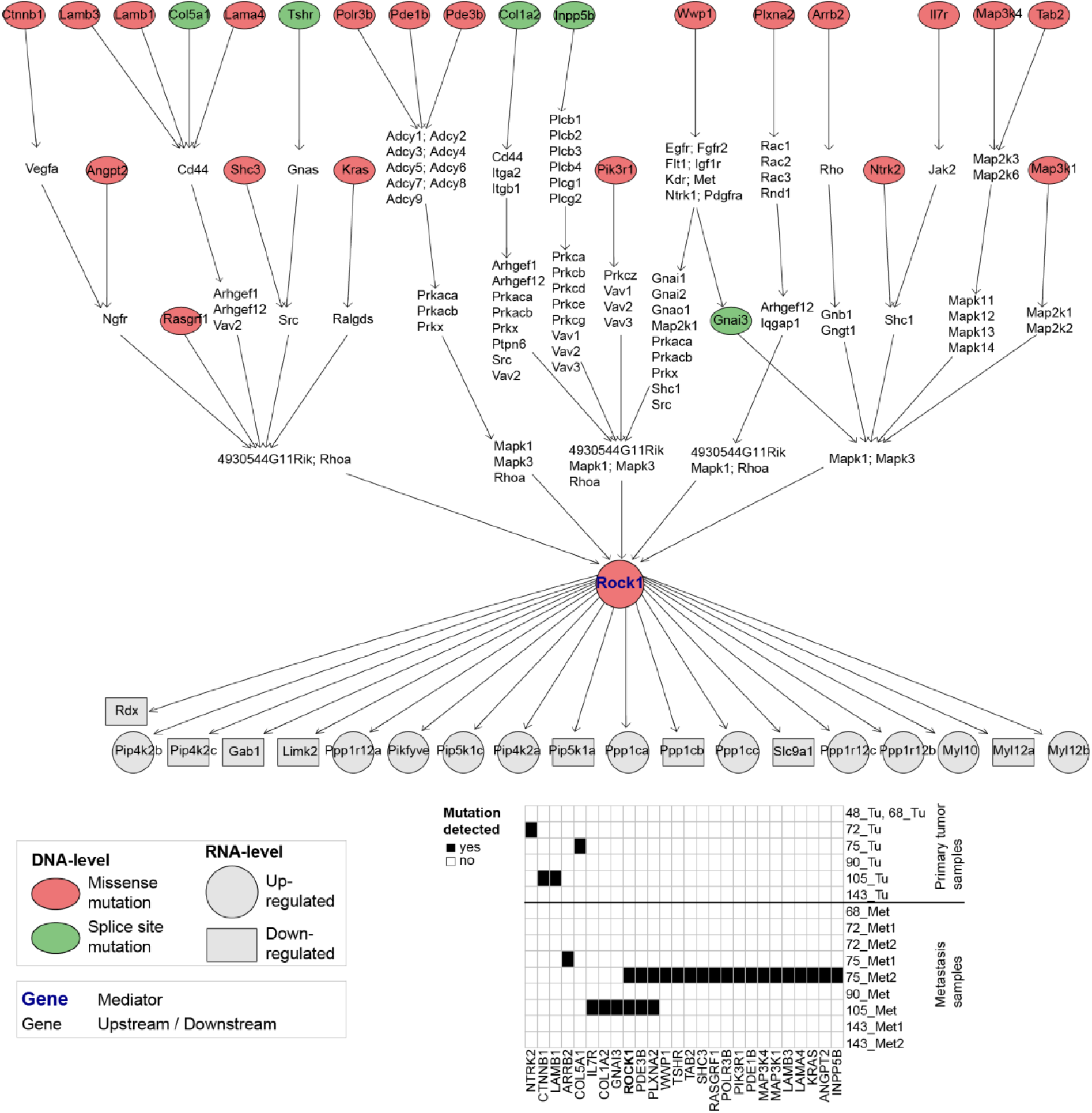
Network of mediator gene *Rock1* in the metastatic versus primary samples comparison. The upstream genes with nonsynonymous genomic variation up to four levels upstream are displayed, as well as the direct downstream genes that are differentially expressed. Also, the genes connecting the upstream mutated genes with the mediator are indicated. In this network, some of the upstream genes are also differentially expressed, but this is not visualized. The table indicates in which samples the mutations were detected.

*Rock1* was overexpressed in the metastasis samples when compared to the primary tumor samples and also in comparison to the normal sample. The overexpression of *ROCK1* has been implicated in metastasis and cell migration^61-65^. In particular, Huang and colleagues demonstrated in a mouse model that the upregulation of *Rock1* is important for metastasis formation in melanoma^63^. In addition, they reported that the activation of the PI3K/AKT pathway can lead to upregulation of *Rock1* in melanoma^63^. Among the upstream mutated genes of *Rock1*, nine genes (*Angpt2, Col1a2, Il7r, Kras, Lamb1, Lamb3, Lama4, Ntrk2, Pik3r1*) are members of the PI3K/AKT signaling pathway^47^. *ROCK1* was also found to be overexpressed and correlated with progression in human bladder cancer^66^, testicular cancer^67^ and human renal cell carcinoma^68^. Lochhead and colleagues reported the detection of activating mutations in *ROCK1* in human cancer that promote tumor progression^69^. In human colorectal cancer, *ROCK1* was significantly upregulated and was associated to poor prognosis^70^.

## Discussion

Our integrative multi-omics analysis of timed melanoma samples identified several interesting aspects of melanoma development. First, we found many mutations in known cancer genes including several melanoma-related genes, as well as genes, such as *Lasp1*, that were previously not reported as prominent in melanoma according to TCGA^26^. *Lasp1* is involved in rearranging the actin cytoskeleton^71^, and *Lasp1* knockout studies indicated that loss of *Lasp1* promotes migration of cells and tumor development^71^. Second, our analysis confirmed that genes involved in remodeling of the extracellular matrix are upregulated in primary melanoma samples, and additionally revealed that this overexpression increases as the tumor progresses over time. Third, the genes *Vegfc* and *Rock1* were identified as important signaling hubs during melanoma progression and metastasis. Fourth, the analysis of genomic heterogeneity between primary tumor samples and metastases revealed that the diversity within mice increases over time even in the absence of extrinsic mutagenic sources such as UV radiation. Fifth, the nonsense-mediated decay process was identified as enriched among the deregulated genes in the metastasis samples when compared to the primary tumor samples and to the normal sample.

We performed network-based integration of genomic and transcriptomic changes to identify important mediator genes that link mutations to downstream gene deregulation. Cancer therapies may be most efficient if they are targeting entire signaling pathways^14,16^. The identification of signaling hubs is a crucial step towards this direction^14^, and our analysis of important mediator genes identified candidate signaling hubs that could be promising points of attack. Among the most striking mediator genes in the analysis of the timed primary samples was the lymphangiogenic growth factor *VEGFC* ^33^, whose direct network revealed many altered genes from the MAPK signaling pathway. *Vegfc* was found to be overexpressed in the primary samples and even more so in the late primary tumor samples. This confirms previous findings that implicate *VEGFC* and lymphatic vessel development in melanoma progression and metastasis^32,34-36^. Additionally, the network integration highlighted several mutated genes upstream of *Vegfc* that may contribute to its overexpression. In particular, each of the late samples showed evidence in the sequence reads for a subset of the *Vegfc* upstream mutations, which demonstrates that these are mutations that occur during the course of melanoma development. Moreover, since this subset of mutations is different between the late samples implying genomic diversity between mice, the network analysis underlined that these genes are all upstream of the same mediator gene *Vegfc* and that genomic diversity may converge to similar transcriptomic changes. Most of the differentially expressed genes in this *Vegfc* network also show stronger deregulation in the later tumor samples, which provides further evidence of sequential increase in MAPK signaling during melanoma evolution.

The analysis of genomic heterogeneity between primary tumor samples and metastases revealed that the diversity within mice increases over time. This increase is significant when considering the time period between 68 and 105 days of tumor evolution. The last mouse, which is from the time point of 143 days after tumor initiation, demonstrated reduced diversity compared to most other mice from earlier time points (Figure 6; Suppl. Figure S5). The other mice from the time points between 68 and 105 days of tumor growth are at most 15 days apart each, whereas the tumor in the very last mouse evolved for another 38 days more than the one before. This could explain the difference in diversity observed here. However, we conjecture that a larger sample size would be necessary to decide whether the very last mouse is an outlier, or whether the diversity is, in fact, reduced after longer time periods of tumor evolution, for instance after more than 140 days.

In general, the increasing diversity over time observed here, though emphasizes that treatment of tumors becomes increasingly difficult, as the tumor progresses over time. Heterogeneous tumors may harbor many different clones that could be resistant to treatment, and could be undetected when using just a single biopsy^72,73^.

The inference of phylogenetic trees resulted in phylogenies that are mostly linear, which is in line with other melanoma studies^56^. We also observe that in most mice, at least one third of the total mutations in the tree are already acquired in the first clone of the tree (Suppl. Figure S6). However, since the average coverage was limited to 65x in our data, the phylogenetic trees should be interpreted with caution. The reliability of the tree inference depends on the depth of coverage^55^.

The differential gene expression analysis of metastasis and primary melanoma samples highlighted the importance of the nonsense-mediated decay process in the metastases, which was significantly overrepresented and upregulated in the metastasis samples. Furthermore, the network-based integration confirmed the prominent role of overexpressed *Rock1* in metastasis formation and cell migration. In addition, the network revealed that many of the upstream mutated genes are members of the PI3K/AKT pathway^47^ and may be contributing to *Rock1* overexpression. Shi et al. have previously reported on the preferential acquisition of mutations upregulating the PI3K/AKT pathway in clinical biopsies of progressive melanomas^49^. Our data confirm these findings in a genetically defined mouse model and suggest that this process of PI3K/AKT upregulation is driven by UV-independent mechanisms. We note that many of the mutations in the *Rock1* network were detected in the hypermutator samples. In the samples without a mutation in the depicted network, *Rock1* upregulation could be caused by epigenetic alterations or other alterations further upstream of *Rock1. ROCK1* is important for organizing the actin cytoskeleton and is therefore involved in contraction, adhesion and migration of cells, and contributes to metastasis and invasion^65,74^. The upregulation of *ROCK1* has been observed in several cancer types^63,74^ and has even been associated with decreased survival^70^. Drugs that target *ROCK1* have been proposed^75,76^, but their effectiveness may also depend on the mutations present in the cells^74,76^. A novel drug that targets *ROCK1* has been recently proposed for hepatoblastoma^77^, which may also be potent in melanoma. Our study adds further evidence that *Rock1* is a promising drug target that could be exploited to reduce the activity of invasive and metastatic cancer cells. It also highlights mutated genes upstream of *Rock1* that could play a role in *Rock1* deregulation.

We note that our study, which relies on the Braf-activated and Pten-deficient mice, focuses on a subset of melanomas. The induced alterations may influence the genomic and transcriptomic changes observed in the tumors. For instance, CDKN2A is commonly lost in human melanoma^26^, but it does not occur in our samples, and could be due to our mouse model^78^. Future studies with different mouse models or human samples are needed to explore the full spectrum of melanoma. However, the joined occurrence of activated Braf and lost Pten is observed in approximately every fifth human melanoma^8^, and therefore highly relevant.

In summary, our comprehensive analysis of melanoma progression and metastasis formation revealed key genes and pathways involved in these processes, which contributes towards an improved understanding of this disease.

## Materials and Methods

### Samples

We employed a mouse model of tamoxifen-inducible Tyr::CreERT2, LSL-*Braf*^V600E^, *Pten*^*fl/fl*^ melanoma^8^ to obtain samples from seven time points of days of tumor growth. The LSL- BrafV600E mice have been received from the Marais Lab^79^. The animal experiments were approved by the Cantonal Veterinary Office Zurich and carried out in accordance to the relevant regulations. Tumor growth was induced on the mouse tail by topical application of 5mg/ml tamoxifen in 100% ethanol. In total we collected seven primary tumor samples and nine samples from metastases in the lymph nodes. Lymph node metastases were not observed in all lymph nodes. The collected metastasis samples were from the inguinal lymph nodes. As a control, a normal tail sample was also collected (Suppl. Table S1).

### Exome analysis

The exomes of the samples were sequenced using the Illumina HiSeq 2500 system to generate 101-basepair paired-end reads. For the computational analysis of the data, the NGS-pipe framework^80^ was customized to include the following steps: Adapters were clipped and low-quality bases trimmed using Trimmomatic^81^. The reads were aligned to the mouse reference genome version mm10 with bwa^82^. Subsequent processing of the alignment files such as merging, PCR duplicate removal and indexing included the use of the SAMtools^83^ and Picard tools^84^ suites. Local realignment around indels and base quality recalibration was done using the Genome Analysis Toolkit (GATK)^85-87^. The depth of coverage was assessed with Qualimap^88^. For the detection of single-nucleotide variants (SNVs), the rank-combination^89^ of deepSNV^90^, JointSNVMix^91^, MuTect^92^, SiNVICT^93^, Strelka^94^, and VarScan2^95^ was employed. Multiple testing correction of the deepSNV p-values was carried out with the R package IHW^96^. Indels were called using the rank-combination of SiNVICT, Strelka, VarDict^97^, and VarScan2. Variants were annotated with the tools SnpSift^98^ and SnpEff^99^. The dbSNP build 142, SNP release v5 from the mouse genome project was used to filter out mutations that are likely not cancerous^100^. Copy number variant (CNV) calling was performed with Excavator^101^ allowing CNVs of length greater or equal than 1Mb, and probability of call greater or equal to 99%. The gonosomes were excluded from the CNV and differential expression analysis since gender differences between the mice may lead to false conclusions. The Variant Effect Predictor^102^ was used to obtain SIFT scores for non-synonymous SNVs in coding regions to determine which mutations are deleterious^23^. The candidate cancer gene database^27^ comprises genes that are potential cancer driver genes in mice. Additionally, it contains the information for each gene whether it is listed in the COSMIC database^103^, and whether it is among the Cancer Gene Census genes^104^. Throughout this study, we referred to this candidate cancer gene database to judge on the importance of genes. In order to get an overview of the functions of specific genes or proteins, the UniProt database was used^105^. The phylogenetic trees were obtained by running Cloe^55^. The parameter for rho, which is the reversion probability, was set to 0.01. Cloe was run 20 different times with different seeds, and then the tree with highest log-posterior probability was selected for visualization and further inspection.

### Transcriptome analysis

RNA-sequencing was done with the Illumina HiSeq 2500 system and 126-basepair single-end reads were generated. Similar as to the exome sequencing analysis, the NGS-pipe framework was employed to analyze the data. The analysis steps included adapter clipping and trimming of low-quality bases using Trimmomatic. Alignment of the reads to the mouse reference genome version mm10 was performed with the STAR aligner^106^. The tool featureCounts^107^ was used to obtain read counts, and differentially expressed genes were called with DESeq2^108^ using a cutoff for adjusted p-values of 0.01. The pathway overrepresentation analysis was performed with WebGestalt^109^ and using all expressed genes as a background gene list that had at least a count of 10 fragments across all samples^110^. The cutoff for the adjusted p-value of an overrepresented pathway was 0.05. Four comparisons were made on the RNA level to detect differentially expressed genes: comparison between all primary tumor samples versus the normal, between late and early primary tumor samples, between metastasis samples and the normal, as well as between the metastasis and primary tumor samples (Suppl. Figure S3).

### Combined analysis

For the computational analysis and visualization of the WES and RNA-seq data, several R packages^111^ were used including ggplot2^112^, reshape2^113^, RColorBrewer^114^, dplyr^115^, gplots^116^, treemapify^117^, VennDiagram^118^, scales^119^, cowplot^120^, ComplexHeatmap^121^, biomaRt^122,123^, OrganismDbi^124^, Mus.musculus^125^, igraph^126^, graph^127^, and Rgraphviz^128^. The criterion for selecting the mediator genes Vegfc and Rock1 was the following: Select those genes from the mediator list that are in the candidate cancer gene database^27^. Among those, choose the gene that is most significant and highly ranked. For Rock1: From the four genes in the list of mediators, Tsc1, Rock1 and Spred1 were in the candidate cancer gene database. Rock1 and Spred1 were highly significant (adjusted p-value < 10^−3^), and Rock1 was higher ranked than Spred1. For Vegfc: The five most highly ranked genes in the list of mediators were all in the candidate cancer gene database. Vegfc was the most significant mediator of the highly ranked genes (adjusted p-value < 10^−3^).

## Author contributions

NB and WK designed the study and acquired the funding. WK, SFM, LF provided the mouse samples. CB prepared and supervised the sequencing. ALM conducted the computational sequencing data analysis and visualization. CD ran the NetICS analysis. ALM inspected and visualized results. ALM drafted the manuscript. All authors read and approved the final manuscript.

## Availability of data and material

The generated WES and RNA-seq data from the 17 samples is made available at the European Nucleotide Archive under accession number PRJEB28651.

## Acknowledgements

The authors would like to thank Hans-Joachim Ruscheweyh for helpful advice concerning the differential gene expression analysis.

