## Supplementary Material for "Dynamic changes of genomic and transcriptomic tumor diversity during melanoma progression"

[†] Deceased 29 August 2018

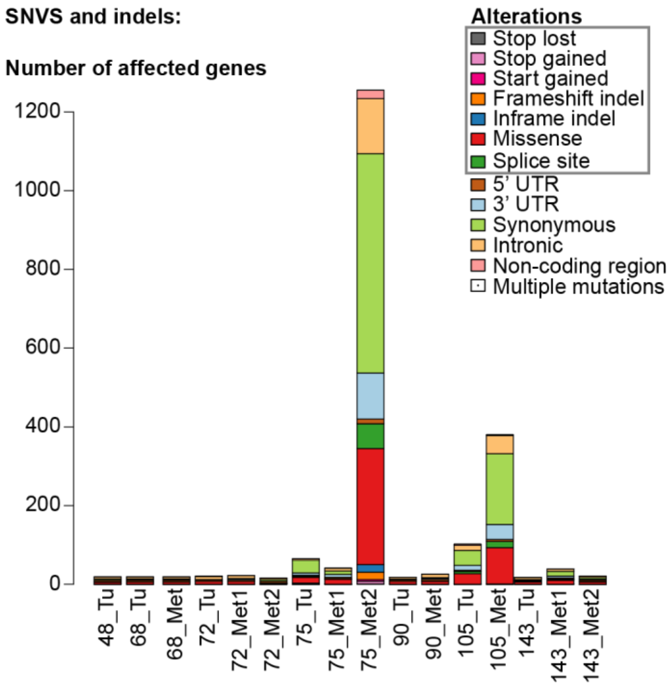


**Supplementary Figure S1.** The numbers of genes that are affected by a SNV or indel per sample. The color indicates the annotation of the mutation. The samples 75_Met2 and 105_Met seem to be hypermutators, since they harbor many more mutations than the other samples.

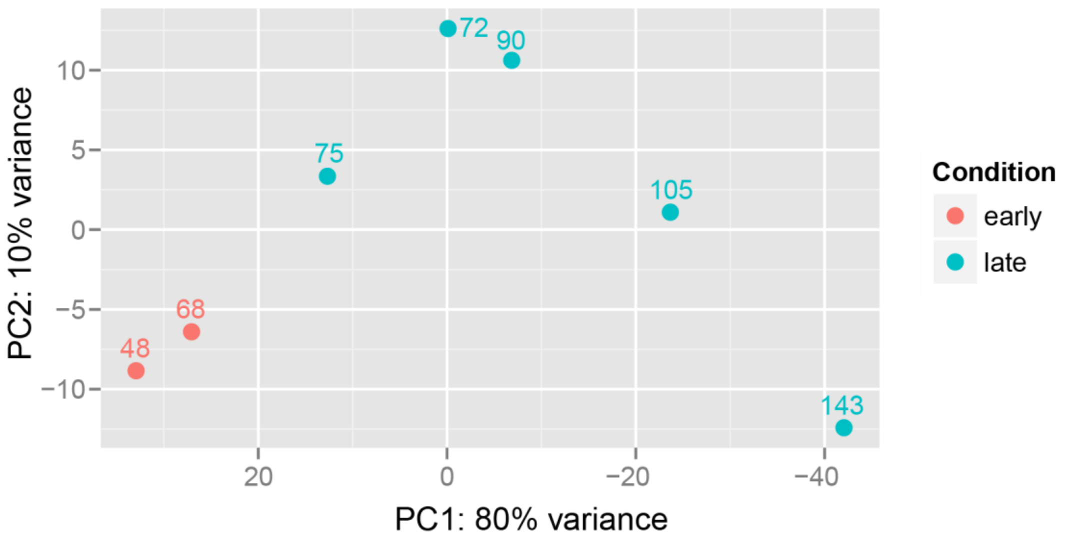


**Supplementary Figure S2.** Transcriptome analysis of timed primary tumor samples. Principal component analysis (PCA) of the expression levels of the 500 most variable genes when including all primary tumor samples. The number of days of tumor growth is indicated in the plot.

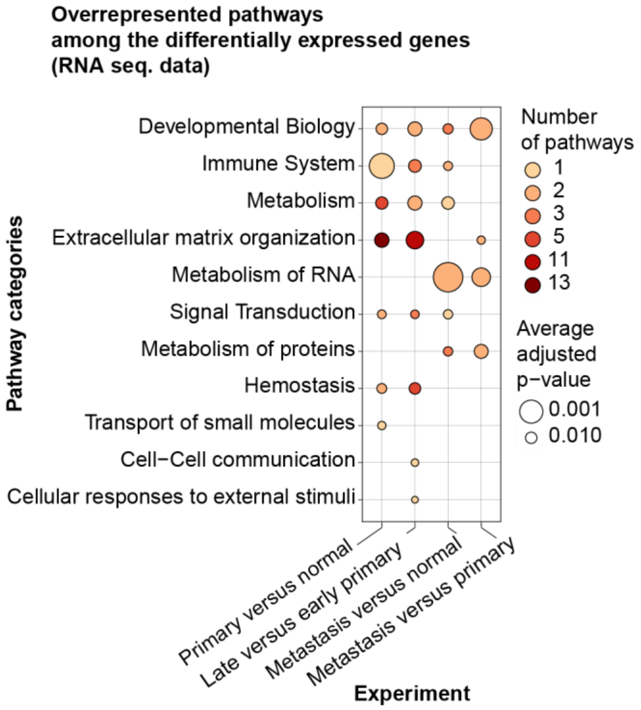


**Supplementary Figure S3.** Overview of the transcriptomic changes. Overrepresentation analysis was performed with the differentially expressed genes using the Reactome pathway data base. The pathways were then assigned to the top-level of the Reactome pathway hierarchy. The number of pathways in each pathway category is indicated. The size of the circle represents the average adjusted p-value of the pathways in each category.

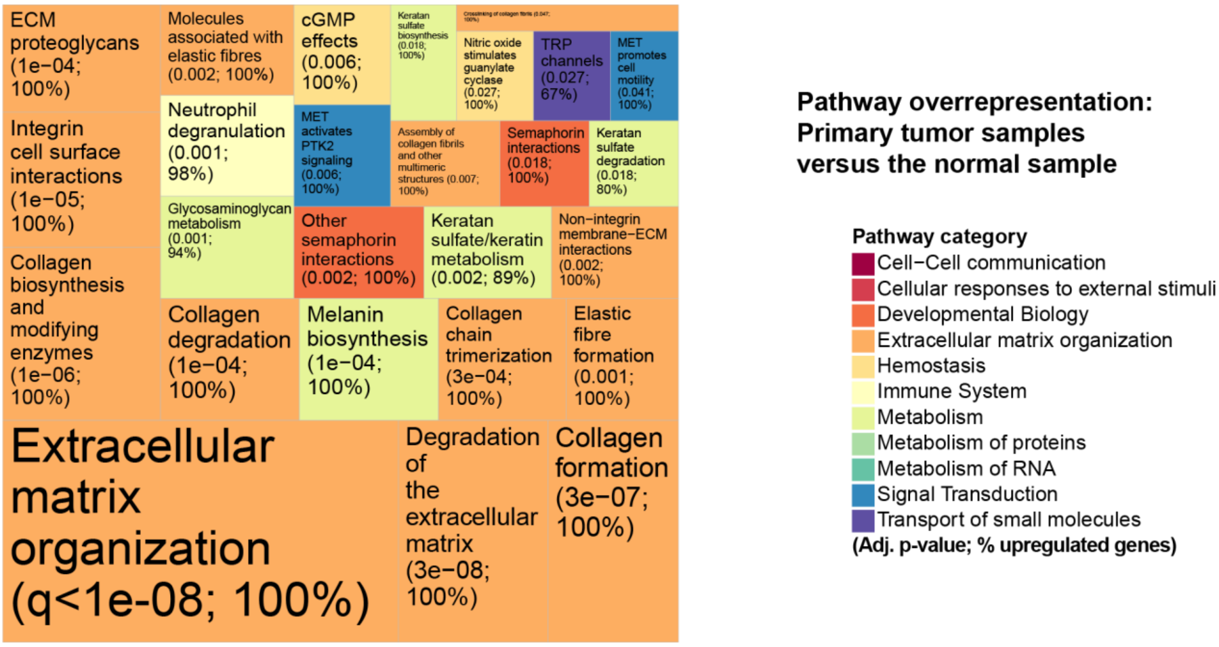


**Supplementary Figure S4.** Pathway overrepresentation analysis was also performed on the differentially expressed genes of the comparison of all primary tumor samples versus the normal sample. The colors represent the top-level pathways of the Reactome hierarchy, and each individual pathway that was significantly overrepresented, is indicated in the treemap. The area of the tiles mirrors the significance level. The percentage of upregulated genes among the deregulated genes in the respective pathway is indicated.

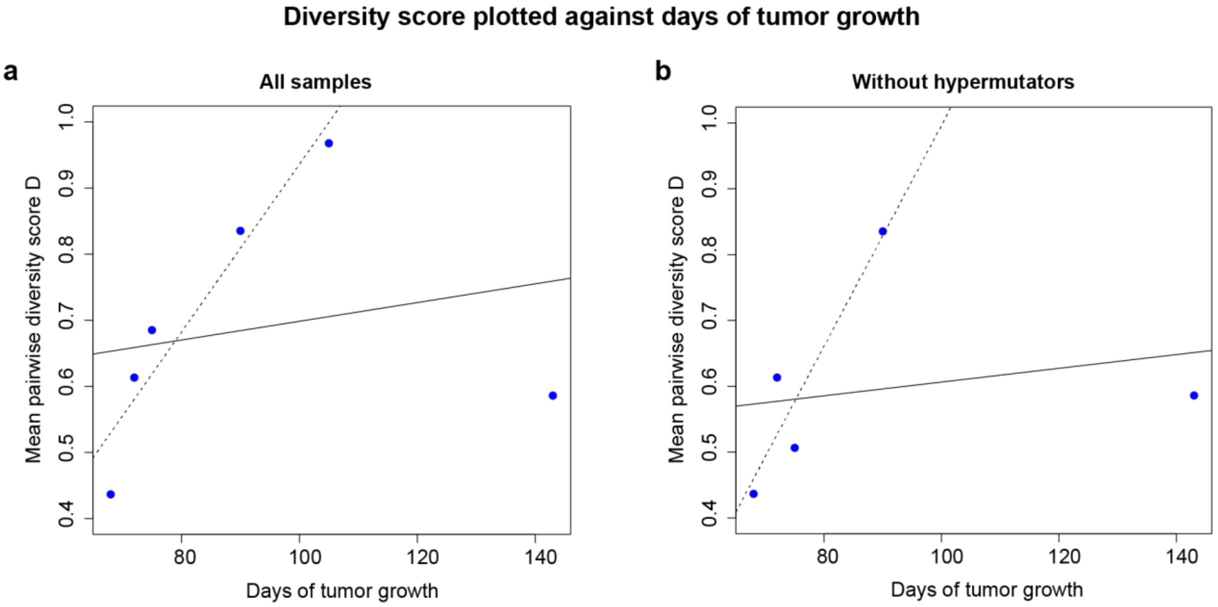


**Supplementary Figure S5.** Diversity score as a function of the numbers of days of tumor growth. The solid regression line in each plot shows an increasing trend. The dashed line is the result of regression without the last mouse at 143 days of tumor growth. The other five mice included here were from the time points 68, 72, 75, 90 and 105 days of tumor growth and are each at most 15 days apart. By contrast, the last mouse is from a much later time point after tumor initiation, i.e. the tumor evolved 38 days more than the second to last mouse at 105 days. Therefore, a separate regression (dashed line) was performed to assess the short term increase in diversity, for the five mice between 68 and 105 days of tumor growth. **(a)** Diversity score plotted against the days of tumor growth while including all samples (solid line slope: 0.0014, p-value = 0.686), and with all but the last mouse at 143 days showing a significant short term increase in diversity (dashed line slope: 0.0127, p-value = 0.013) **(b)** Diversity score plotted against the days of tumor growth while including all samples but the two hypermutators 75_Met2 and 105_Met (solid line slope: 0.001; p-value = 0.729). Without the two hypermutators, the observation at 105 days of tumor growth cannot be included any more, since only the primary tumor sample of that mouse is left. Therefore, five observations remain (68, 72, 75, 90, 143 days of tumor growth). The regression without the last mouse at 143 days of tumor growth also shows a significant short term increase in diversity (dashed line slope: 0.0168, p-value = 0.074).

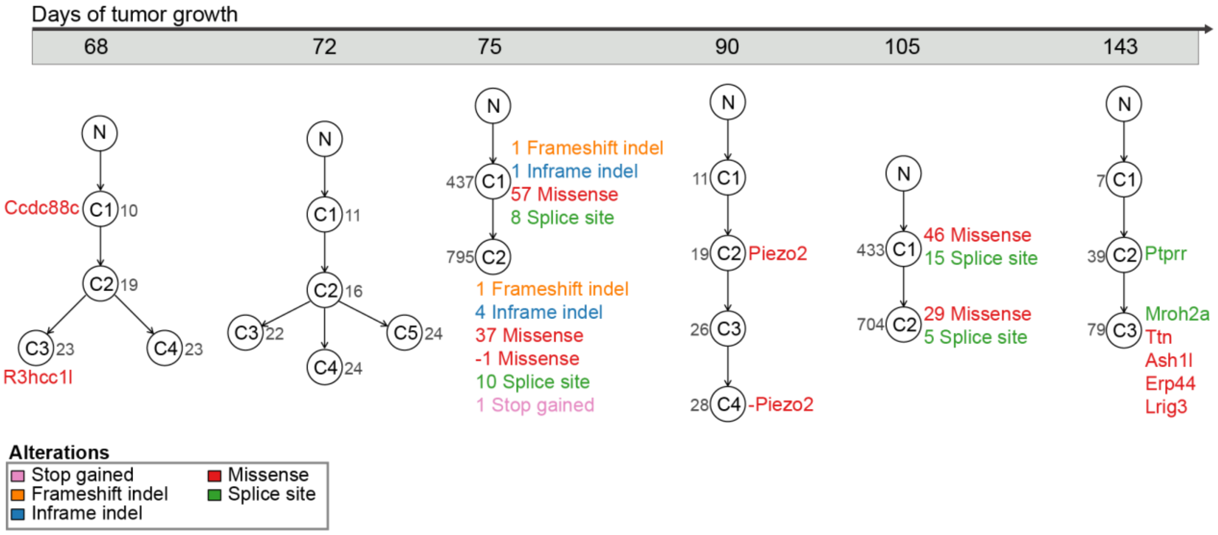


**Supplementary Figure S6.** The phylogenetic trees as obtained by running Cloe[52]. Cloe was run 20 different times with different seeds. Subsequently, the tree with the highest log-posterior probability was selected for each mouse. The highlighted genes are those that are in the candidate cancer gene database and harbor a non-silent mutation in a coding region. The mouse at 75 days of tumor growth was the one with the hypermutator metastasis. Altogether in primary tumor and both metastasis samples, there were 2,622 mutations, which is too many as input for Cloe. Therefore, 800 mutations were subsampled in each of the 20 Cloe runs for the mouse at 75 days of tumor growth. The grey numbers next to the clones indicate the total number of mutations that were assigned to the respective clone. We observe that for the first five mice (68, 72, 75, 90, 105 days of tumor growth), at least one third of the total mutations in the tree are already acquired in the first clone of the tree. By contrast, in the last mouse, only 9% of all mutations in the tree occur already in the first clone.

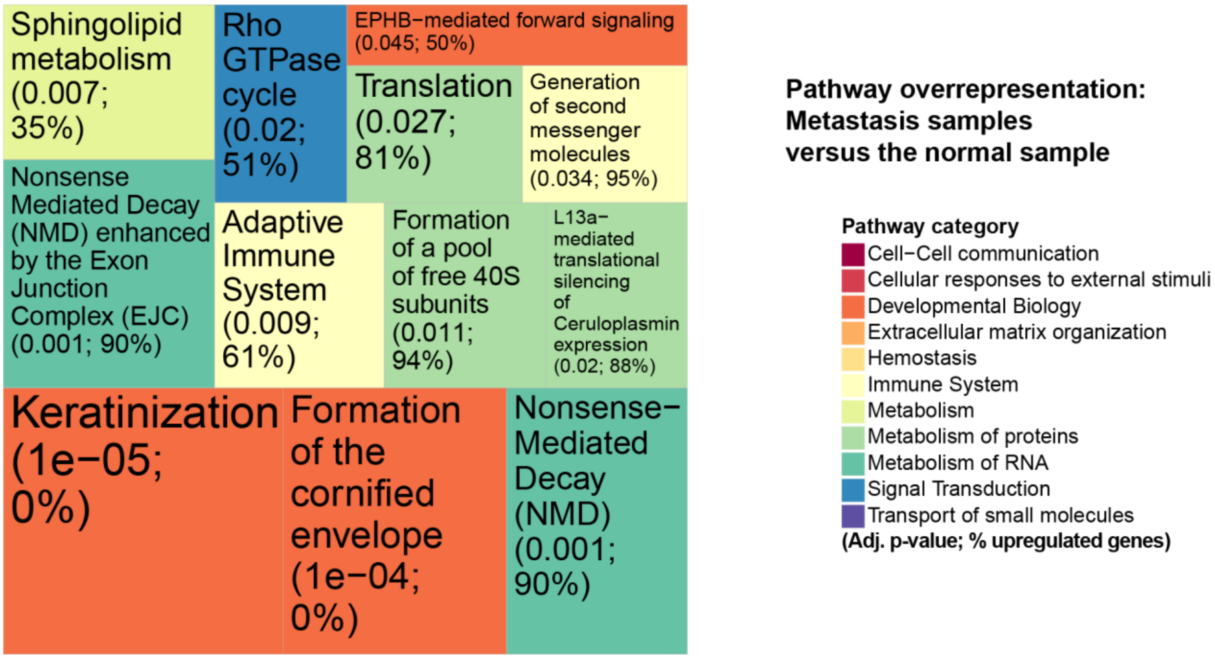


**Supplementary Figure S7.** Pathway overrepresentation analysis was also performed on the differentially expressed genes of the comparison of all metastasis samples versus the normal sample. The colors represent the top-level pathways of the Reactome hierarchy, and each individual pathway which was significantly overrepresented, is indicated in the treemap. The area of the tiles mirrors the significance level. The percentage of upregulated genes among the deregulated genes in the respective pathway is indicated.

**Supplementary Table S1.** Overview of all 17 samples.

| Mouse # | Sample ID | Sample Type | Days of Tumor Growth | Average Coverage in WES sample |
|---|---|---|---|---|
| 1 | 48_Tu | primary tumor | 48 | 67x |
| 2 | 68_Tu | primary tumor | 68 | 74x |
| 2 | 68_Met | metastasis | 68 | 66x |
| 3 | 72_Tu | primary tumor | 72 | 67x |
| 3 | 72_ Met1 | metastasis | 72 | 73x |
| 3 | 72_ Met2 | metastasis | 72 | 63x |
| 4 | 75_Tu | primary tumor | 75 | 40x |
| 4 | 75_ Met1 | metastasis | 75 | 77x |
| 4 | 75_ Met2 | metastasis | 75 | 39x |
| 5 | 90_Tu | primary tumor | 90 | 57x |
| 5 | 90_Met | metastasis | 90 | 63x |
| 6 | 105_Tu | primary tumor | 105 | 79x |
| 6 | 105_Met | metastasis | 105 | 67x |
| 7 | 143_Tu | primary tumor | 143 | 47x |
| 7 | 143_Met1 | metastasis | 143 | 69x |
| 7 | 143_Met2 | metastasis | 143 | 79x |
| 8 | normal | Normal tail sample | 0 | 73x |

The primary tumor samples and the normal sample are from the tail. The metastases are from the lymph nodes.

**Supplementary Table S2.** Mediator genes from the NetICS analysis[15] including the deregulated genes from the "primary versus normal" analysis, as well as all mutations in the primary tumor samples.

| Gene Name | Adjusted p-value | Rank by proximity to altered genes |
|---|---|---|
| **STAT3** | 0.0184 | 1 |
| **VEGFC** | $< 10^{-3}$ | 2 |
| **STAT1** | 0.0184 | 3 |
| **TYK2** | 0.044872 | 5 |
| **SOCS2** | 0.010243 | 6 |
| **SHC3** | $< 10^{-3}$ | 7 |
| **SHC4** | $< 10^{-3}$ | 8 |
| **TYR** | $< 10^{-3}$ | 9 |
| **STAT5A** | 0.025049 | 10 |
| **STAT5B** | 0.0184 | 11 |
| **VEGFD** | 0.010243 | 15 |
| **PREX1** | $< 10^{-3}$ | 19 |
| **PIK3R3** | $< 10^{-3}$ | 20 |
| **TH** | $< 10^{-3}$ | 21 |
| **STAT6** | 0.04968 | 22 |
| **PIP5K1B** | $< 10^{-3}$ | 26 |
| **WASF1** | $< 10^{-3}$ | 34 |
| **SCN8A** | $< 10^{-3}$ | 41 |
| **TLR4** | 0.036529 | 42 |
| **PI4K2B** | $< 10^{-3}$ | 46 |
| **VCAN** | 0.010243 | 51 |
| **PGF** | $< 10^{-3}$ | 56 |
| **RRM2B** | 0.040833 | 65 |
| **RRM2** | 0.040833 | 66 |

| | | |
|---|---|---|
| **PPARGC1A** | < $10^{-3}$ | 68 |
| **TYROBP** | < $10^{-3}$ | 69 |
| **XDH** | < $10^{-3}$ | 80 |
| **PDE1C** | < $10^{-3}$ | 82 |
| **PDE1B** | < $10^{-3}$ | 83 |
| **PDE3A** | < $10^{-3}$ | 84 |
| **MYC** | 0.044872 | 87 |
| **PDGFRA** | 0.031543 | 88 |
| **TREM2** | < $10^{-3}$ | 89 |
| **PTPRC** | 0.040833 | 91 |
| **PDGFRB** | 0.0184 | 93 |
| **PDE7B** | < $10^{-3}$ | 98 |
| **PDE4A** | 0.010243 | 99 |
| **PDE10A** | < $10^{-3}$ | 100 |
| **PDE11A** | 0.010243 | 101 |
| **PDE5A** | < $10^{-3}$ | 106 |
| **NMNAT2** | < $10^{-3}$ | 108 |
| **SOX4** | 0.010243 | 113 |

The rank is according to the proximity to upstream mutated genes and downstream differentially expressed genes. These mediator genes can be regarded as signaling hubs that possibly transfer the abnormal signaling from mutated upstream genes and lead to differential expression of downstream genes. The p-value is from a permutation test that permutes mutated and differentially expressed genes.

**Supplementary Table S3.** Mediator genes from the NetICS analysis[15] including the deregulated genes from the “metastasis versus primary samples” analysis, as well as mutated genes detected in either the primary tumor samples or the metastases.

| Gene Name | Adjusted p-value | Rank by proximity to altered genes |
|---|---|---|
| **SHC3** | $< 10^{-3}$ | 20 |
| **TSC1** | 0.04968 | 22 |
| **ROCK1** | $< 10^{-3}$ | 40 |
| **SPRED1** | $< 10^{-3}$ | 50 |

The rank is according to the proximity to upstream mutated genes and downstream differentially expressed genes. These mediator genes can be regarded as signaling hubs that possibly transfer the abnormal signaling from mutated upstream genes and lead to differential expression of downstream genes. The p-value is from a permutation test that permutes mutated and differentially expressed genes.
